# Wound-like tumor periphery in human breast cancer predicts a convergent drug nonresponse

**DOI:** 10.1101/2021.11.02.467008

**Authors:** Lianhuang Li, Xiaoxia Liao, Fangmeng Fu, Gangqin Xi, Deyong Kang, Jiajia He, Wenhui Guo, Lida Qiu, Zhonghua Han, Xingfu Wang, Qingyuan Zhan, Sixian You, Jianxin Chen, Chuan Wang, Stephen A. Boppart, Haohua Tu

## Abstract

A significant portion of breast cancer patients are nonresponsive to well-established drugs and destined for a poor outcome regardless of molecular subtype. Although several (multiparameter) molecular markers have predicted their resistance to some of these drugs, profound uniparameter markers predictive of a convergent nonresponse to all these drugs remain elusive. We employ co-registered standard-multiphoton histology to representatively sample a few peripheral niches of the primary tumor, so that hundreds of patients can be stratified with either a wound-like or non-wound tumor periphery. With no fitting variable, this simple uniparameter morphological marker is: (a) highly sensitive and specific to predict a multidrug-nonresponsive phenotype that accounts for the majority of recurrence or death, independent of the molecular subtype or related adjuvant drug selection, clinical endpoint (disease-free versus overall survival), and hosting medical center; (b) robust against intratumor heterogeneity and valid at the earliest clinicopathological stage; and (c) dominant in predicting prognosis in the context of routine clinicopathological markers. Considering the mechanistic link between a wound-like extracellular matrix and a microenvironment supporting migratory or mesenchymal tumor cells, we attribute these unusual capabilities to an epithelial-mesenchymal transition nature of the morphological marker long sought after by pathologists.

## Introduction

The molecular portraits of human breast cancer (1) have not only inspired similar molecular characterizations of other cancer types but also justified the molecular subtype-classified drug selection (2) and treatment guideline (3). As part of precision medicine, the corresponding resistance to various first-line drugs has also been classified by the molecular subtype into: (i) chemoresistance (to anthracycline-taxane) in the chemotherapy of triple-negative breast cancer (TNBC) (4); (ii) endocrine resistance (to tamoxifen) in the hormone therapy of human epidermal growth factor receptor 2 negative luminal (HER2-&luminal) breast cancer (5); and (iii) large molecule resistance (to trastuzumab) in the targeted therapy of HER2+ breast cancer (6). These diverse forms of drug resistance have been typically inferred from *in vitro* cell/tissue culture (6, 7), *in vivo* animal models (8), and the pathological complete response (pCR) of human patients in the neoadjuvant setting often correlated with the poor prognosis of molecular subtype-classified patient cohorts (9). Despite the identification of some markers or pathways for the chemo-, endocrine, and large molecule resistance (4-9), it remains ambiguous whether most drug-resistant patients with a poor outcome differ significantly among the three treatment subgroups according to some molecular subtype-dependent resistant mechanisms or they share a convergent resistant marker and underlying mechanism independent of the molecular subtype (10). The demonstration of the latter, particularly in the form of a simple uniparameter marker (11), would be valuable to identify more profound targets for new anti-cancer drugs and high-risk patients that warrant alternative unconventional therapies (12). With an emphasis on human tumor microenvironment, we aim to demonstrate the latter in a large-scale clinical study without the traditional clinical translation from cells/animals to humans.

Due to the complexity of cancer drug resistance, three major efforts have been taken to design an “idealized” study in order to proactively avoid plausible confounding factors. First, we choose the adjuvant over neoadjuvant setting for systemic treatment of nonmetastatic patients, even though the latter is gaining popularity to study drug resistance (13). Thus, we use disease-free survival (DFS) and overall survival (OS) as the endpoints to assess the *drug nonresponse* (or treatment failure) at occult metastatic sites (11), rather than pCR (as a surrogate endpoint of DFS/OS) to assess the *drug resistance* in the primary tumor (9). The differentiation between these two often interchangeable terms avoids the confounding factor of pCR as a paradoxical (14) or weak predictor of DFS/OS (15). Second, we require all patients to be diagnosed by symptom rather than (mammographic) screening (Table S1), so that the plausible confounding factor of overdiagnosis is eliminated (16). Also, our symptom-detected patients avoid the overrepresentation of luminal patients with advanced ages (Table S2), mitigating another confounding factor on DFS/OS that these patients experience higher mortality from nonmalignant causes and necessitate specific models for correction (17). Third, we demand all recruited patients to be treated with chemotherapy regardless of the molecular subtype, despite the availability of gene expression assay (18) or computational risk assessment (19) to spare many luminal patients with endocrine therapy from this additional non-first-line therapy, which may be nonbeneficial and even harmful. Plausibly low absolute benefit of the chemotherapy is countered by the absence of the confounding factor that similar relative benefit of chemotherapy exists in all molecular subtype-classified subgroups (20). It should be noted that symptom-detected patients with relatively young ages are rather tolerable to the side effects of chemotherapy.

It would be challenging in a high-income country to recruit representative patients at multiple medical centers by simultaneously avoiding the neoadjuvant treatment, screening mammography, gene expression assay, and computational risk assessment, all of which have improved the standard care for nonmetastatic breast cancer. In contrast, the period of this retrospective study (2003-2014) in China offers a unique window to perform our “idealized” study without these advanced technologies and related confounding factors, due to the lack of technical expertise to assess pCR, a national program for breast cancer screening, and the logistics to support the gene expression assay or computational risk assessment (21). Although these deficiencies were suboptimal for patient care, they put us in a good position to identify the representative drug-nonresponsive patients with “full” treatment (Fig. 1a), except for the tendency of HER2 patients to forgo the trastuzumab-based targeted therapy with uncertain cost-to-benefit ratio (22). Indeed, we find that our TNBC, HER2-&luminal, and HER2+ patients share a convergent drug-nonresponsive phenotype with a common uniparameter marker of wound-like tumor periphery (23), which predicts most of the observed recurrence or death events with high accuracy. Also, we demonstrate the robustness of this morphological marker against the intratumor heterogeneity (24) that may reduce the clinical utility of many molecular markers (25). Finally, from a rare perspective of histology-like imaging, we provide evidence on an epithelial-mesenchymal transition (EMT) nature of this marker (26).

**Figure 1.**
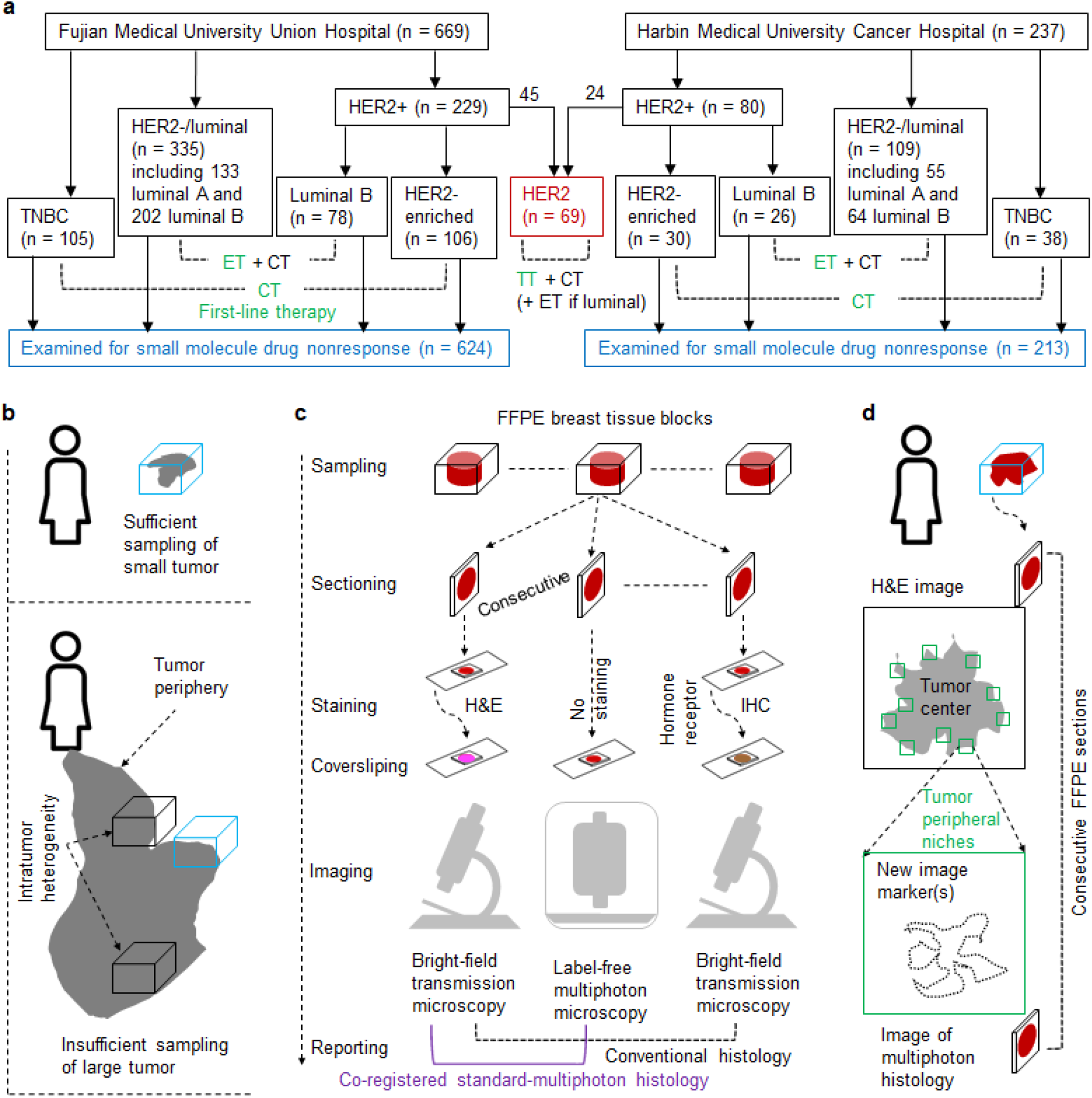
Patients and methods. (a) Included patients from two medical centers undergoing molecular subtype-classified therapies (CT, ET, and TT); (b) Sufficient sampling of a small (≤2 cm) tumor in one patient by one FFPE block (top), and insufficient sampling of a large (∼5 cm) tumor in another patient by multiple FFPE blocks, which were used for H&E histology but only one of them (blue box) that intersected the tumor periphery was selected for the multiphoton histology (bottom). (c) Inclusion of multiphoton histology into routine formalin-fixed paraffin-embedded (FFPE) workflow of breast cancer H&E histology and immunohistochemistry (IHC). (d) Co-registered standard-multiphoton histology to co-register a large-scale H&E image of one FFPE section with small-scale high-resolution images of multiphoton imaging at multiple niches (regions of interest) at the tumor periphery.

## Materials and Methods

### Patients with clinicopathological and treatment information

Under a protocol approved by the Institutional Review Boards of Fujian Medical University Union Hospital (or Harbin Medical University Cancer Hospital), we collected anonymous data from 669 (or 237) female invasive breast cancer patients underwent primary tumor surgery within 3 months of initial diagnosis between 2003 and 2014, after excluding a small portion (26%) of patients according to: (a) presence of neoadjuvant therapy; (b) no symptom at diagnosis; (c) absence of chemotherapy; and (d) other factors such as metastasis at diagnosis, ductal carcinoma *in situ*, missing clinicopathological information or DFS follow-up. The clinicopathological information (tumor size, lymph node status, histological grade, and age) were obtained according to the classical TNM staging and the Nottingham histologic grade, while DFS was defined to be the time from the date of diagnosis to that of recurrence/death or ended follow-up if no recurrence/death occurred. For the 669 patients from Fujian Medical University Union Hospital, we also collected OS data defined as the time from the date of diagnosis to that of death or ended follow-up if the patient survived (Tables S1, S2). Depending on the molecular subtype detailed below, the included patients were classified into different subgroups and went through various first-line therapies (anthracycline-taxane-based chemotherapy CT, tamoxifen-based endocrine therapy ET, or trastuzumab-based targeted therapy TT) and possible non-first-line therapies (2) (Fig. 1a).

### Molecular subtype from immunohistochemistry

The molecular subtype of each patient was determined by the expression of estrogen receptor (ER), progesterone receptor (PR), HER2, and Ki67 via immunohistochemistry. Tumors with HER2 staining scores of 0 or 1+ (3+) were defined as HER2-(HER2+). For HER2 staining scores of 2+, the unamplified (amplified) results of *in situ* hybridization were defined as HER2-(HER2+). A patient was classified into one of three major molecular subtypes: (i) TNBC (HER2-, ER-, and PR-); (ii) HER2+&luminal (HER2-, ER/PR+), which can be subdivided into luminal A (HER2-, ER/PR+, and Ki67-low) and HER2-&luminal B (HER2-, ER/PR+, and Ki67-high); and (iii) HER2+, which can be subdivided into HER2+&luminal B (HER2+, ER/PR+) and HER2-enriched (HER2+, ER-, and PR-) (Table S1).

### Sample preparation and subsequent histology

One formalin-fixed paraffin-embedded (FFPE) tissue block (∼2 cm in size) containing both the primary tumor and adjacent normal appearing tissue was selected from archived blocks for each patient (Fig. 1b). A microtome cut two consecutive serial sections (5 μm) from the tissue block. One section was deparaffinized by alcohol & xylene and stained with H&E staining for standard histology (whole slide imaging), and the other section was simply deparaffinized for label-free multiphoton histology (Figs. 1c, 1d). The standard histology was conducted by a commercial whole slide scanner (VM1000, Motic, China) at a resolution of ∼0.5 μm using a 40× microscope objective. The multiphoton histology was performed by a commercial microscope equipped with a Ti:Sapphire laser emitting 810-nm 150-fs pulses (LSM 880, Zeiss, Germany). The back-reflected signals were simultaneously obtained via two spectral channels of second harmonic generation SHG (395-415 nm) and two-photon auto-fluorescence TPAF (428-695 nm). A 20× microscope objective (NA 0.8, Zeiss, Germany) was employed to acquire images at a resolution of 0.8 μm and a FOV of 0.5×0.5 mm^2^. Larger scale imaging of 2.8×2.8 mm^2^ was enabled by a computer controlled mechanical stage through automatic mosaicking. The average standard (or multiphoton) histology time on one section/patient was 15 minutes (or 70 minutes).

### Co-registered standard-multiphoton histology

We co-registered label-free dual-channel (SHG-TPAF) multiphoton microscopy/histology with standard (H&E) histology. The method consists of three steps (Figs. 1c, 1d): (i) a histotechnologist performed serial sectioning (5 μm) to retrieve one section, stained it with H&E, and conducted whole slide imaging via bright-field transmission microscopy; (ii) from the whole slide H&E image, a pathologist isolated tumor periphery (i.e. invasive front/margin) from adjacent tumor center and normal appearing tissue by alternating low- and high-magnifications, and randomly marked 5-15 niches (2.8-mm-sized regions of interest) to sufficiently sample the tumor periphery; and (iii) an imaging scientist retrieved an adjacent section from the same serial sectioning, deparaffined this section on a microscope slide to avoid plausible undesirable effects of paraffin on subsequent imaging, scanned the deparaffined section by the multiphoton histology at low magnification to enable co-registration with the whole slide H&E image, and repeated the same multiphoton histology at a higher magnification for the marked niches. The standard and multiphoton histology images were only co-registered at the higher magnification. This method synergistically integrated the standard histology to locate the tumor periphery at a coarse scale with the multiphoton histology to recognize new morphological markers at a finer scale.

### Determination of tumor-periphery morphology

Three inspectors (L. Li, G. Xi, and J. He), who were blind to the OS and DFS data, independently examined the morphological state of all probed niches in each FFPE section (patient) from the SHG-TPAF images. For a given tumor peripheral niche, its morphology was classified as “non-wound”, “wound-like”, or “intermediate” (see definition below). This morphological state was detected at the niche-scale of 2.8 mm and an optical resolution of 0.8 µm throughout this study. The prevalence of the “intermediate” morphology was relatively low as most (>60%) niches were conclusively classified into either the “wound-like” or “non-wound” morphology. Each niche was then assigned a score of “-1/non-wound”, “+1/wound-like”, or “0/ intermediate”. The wound score of the corresponding tumor/patient was then determined by averaging the scores over all niches and inspectors. In this way, wound-like tumor periphery (WTP) of a given patient was determined as WTP1 (wound score <0), WTP2 (wound score =0), or WTP3 (wound score >0). Thus, all patients under study could be stratified into three categories of WTP1-3 (Table S3), or two categories of WTP-(WTP1/WTP2) and WTP+ (WTP3) (Tables S4-S6). The typical speed of a trained inspector was ∼10 minutes/section. Inter-observer discordance of the above scoring was evaluated by intraclass correlation coefficient. The high inter-observer agreement on WTP-/WTP+ determination at patient-level was validated by an intraclass correlation coefficient value of 0.87 among the three inspectors.

## Results

### Two competing wound-related morphologies of tumor peripherical niches

Our multiphoton histology revealed a small (∼1.5 mm) tumor in an exploratory study of freshly excised human breast tumors. Interestingly, this tumor consisted of one half with a “wound-like” morphology characterized by multidirectionally aligned SHG-highlighted fibrillar collagen (presumably collagen Type I) and abundant 2PAF-highlighted vessels that supported the migration of mesenchymal tumor cells, and the other half with a “non-wound” morphology (23) characterized by SHG-highlighted layered collagen (presumably collagen Type IV) and 2PAF-highlighted tumor mass that formed a discernable tumor boundary and supported the proliferation of epithelial tumor cells (Fig. 2a). We hypothesized that the two extreme or opposite morphologies would compete in tumor progression, and the periphery (invasion front/margin) of the subsequently enlarged tumor could be representatively probed/imaged by a few 2.8-mm-squared niches or tumor microenvironments (Fig. 1d). Each of the niches would then be assigned a “wound-like”, “non-wound”, or “intermediate” morphology.

**Figure 2.**
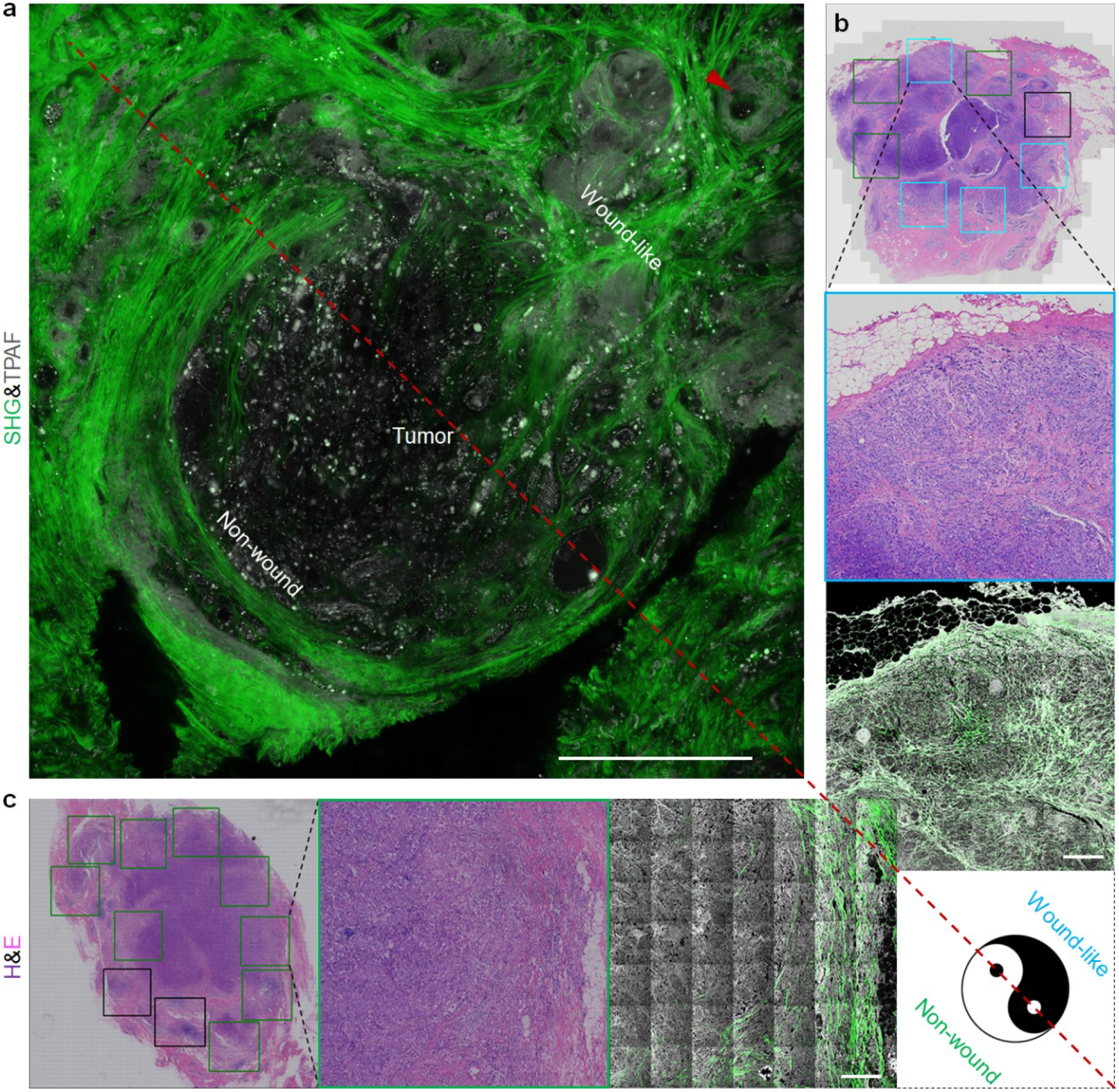
Concept of competing “wound-like” versus “non-wound” tumor periphery (lower right). (a) “Wound-like” tumor periphery (WTP) characterized by SHG-highlighted sparse collagen fibers randomly aligned among TPAF-highlighted angiogenic vessels (arrowhead) and discontinuous tumor cells/clusters without a clear tumor boundary, and “non-wound” tumor periphery characterized by a discernable tumor boundary between TPAF-highlighted continuous tumor mass with no/little fibrillar collagen and surrounding SHG-highlighted collagen in a layer-like formation. (b) Low-vs. high-power H&E images of one patient (blue-squared niche in the large-scale H&E image) and co-registered SHG-TPAF image with the “wound-like” tumor periphery. (c) Similar H&E images from another patient (green squared niche in the large-scale H&E image) and co-registered SHG-TPAF image with the “non-wound” tumor periphery. A patient is WTP+ (WTP-) if there are more (less) blue squared niches than green squared niches, while the black squared niches (“intermediate” tumor periphery) in the low-power H&E images are ignored. Scale bar: 400 µm.

We then performed the retrospective study based on archived FFPE blocks. Personized WTP was determined by the dominancy of “wound-like” or “non-wound” morphology via the niche-scale SHG&TPAF images co-registered with the section-scale (∼2 cm) H&E image (see Methods), despite the wide variation of observed “non-wound”, “wound-like”, and “intermediate” morphologies (see examples in Figs. S1-S3). As a result, one patient with a dominant “wound-like” tumor periphery (WTP+) had a “wound-like” tumor peripheral niche in the FFPE section that retained the similar SHG-highlighted collagen distribution to that of the fresh tissue, even though the TPAF-highlighted vessels collapsed and were less obvious (Fig. 2b). This patient was easily differentiated from a WTP-patient with a dominant “non-wound” tumor periphery (Fig. 2c).

### Convergent phenotype nonresponsive to molecular subtype-classified therapies

It is intuitively to link WTP+ (or WTP-) with high (or low) metastatic risk of the “wound-like” (or “non-wound”) tumor periphery supporting mesenchymal (or epithelial) tumor cells. To examine this intuition, we tested this marker on the 105 CT-treated TNBC patients from Fujian Medical University Union Hospital (Fig. 1a) and identified 27 of 43 patients with <5-year DFS against 4 false positives (Fig. 3a), i.e. attained 63% sensitivity and 94% specificity (*p* = 1×10^−10^) (Table S4). Thus, WTP+ patients accurately accounted for most of the anthracycline-taxane-nonresponsive patients. Similarly, for the 335 ET-treated HER2-&luminal patients from this hospital (Fig. 1a), we identified 70 of 101 patients with <5-year DFS against 57 false positives (Fig. 3a), i.e. attained 70% sensitivity and 76% specificity (*p* = 2×10^−14^) (Table S4). Again, WTP+ patients accurately accounted for most of the tamoxifen-nonresponsive patients, as a deceased specificity in comparison to the TNBC subgroup was compensated by an increased sensitivity. Together, we observed a common breast cancer phenotype nonresponsive to two distinct molecular subtype-classified therapies or drugs, which contrasted each other with a low-sensitivity high-specificity (LSHS) profile from one molecular subtype versus a high-sensitivity low-specificity (HSLS) profile from another molecular subtype (Fig. 3a). In other words, the TNBC and HER2-&luminal patients shared a common drug-nonresponsive phenotype responsive for most <5-year recurrences or deaths.

**Figure 3.**
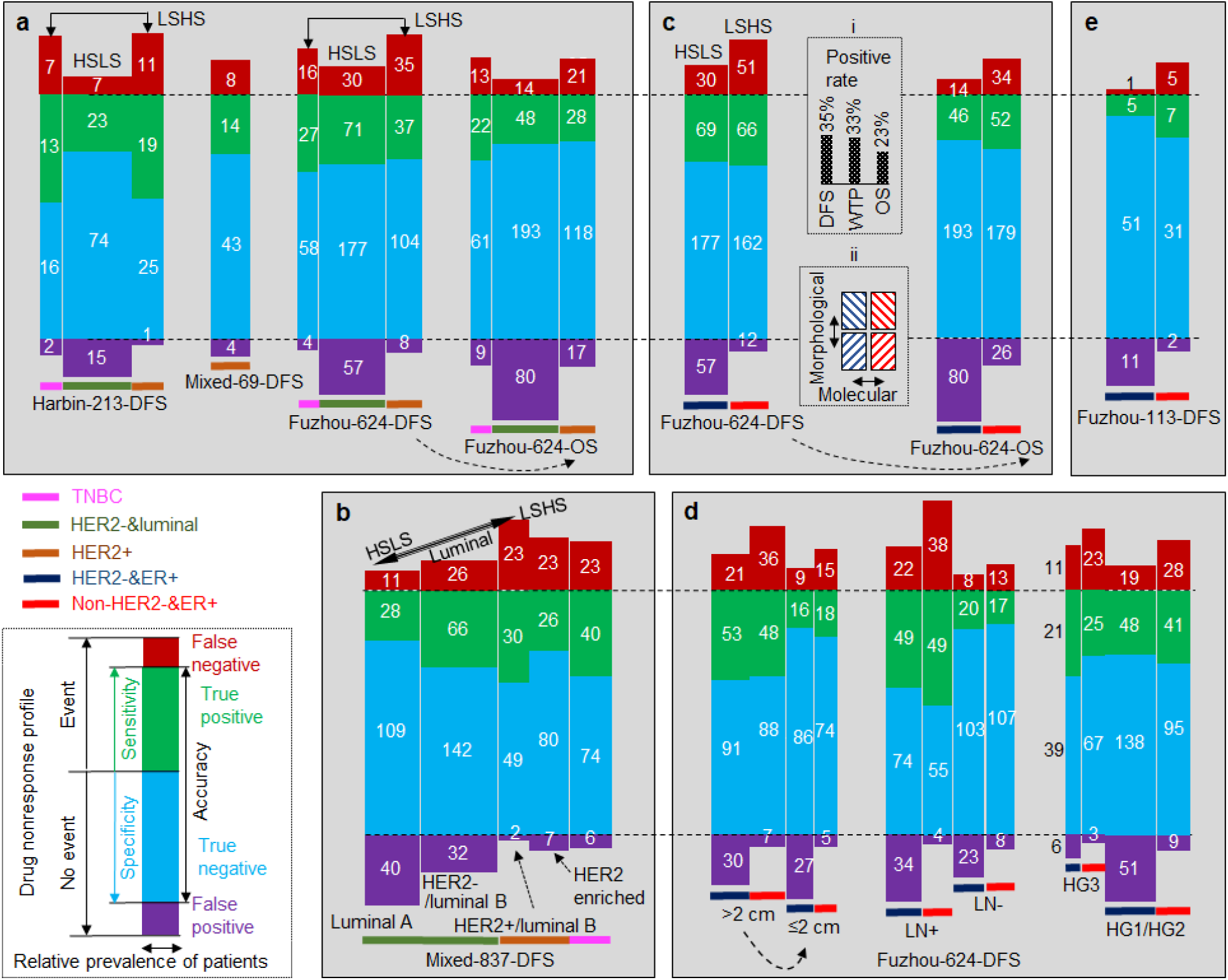
Drug nonresponse profiles (DNP) of distinct patient subgroups from WTP prediction of <5-year DFS/OS. (a) molecular subtype-dependent DNPs observed from 624 patients of Fujian Medical University Union Hospital, 213 patients of Harbin Medical University Cancer Hospital, and 69 patients of both hospitals; (b) more subtle molecular subtype-dependent DNPs observed from 837 patients of both hospitals; (c) Simplified DNP with HSLS vs. LSHS dual-profile pattern for the 624 patients with either DFS or OS endpoint. see text for details on two insets; (d) DNP dual-profile pattern of the 624 patients robust against two categories of tumor size, lymph node (LN) status, and histological grade (HG); and (e) Validity of DNP dual-profile pattern for 113 clinicopathologically low-risk patients among the 624 patients.

Remarkably, the same dual-profile drug nonresponse was reproduced by 38 TNBC and 119 HER2-&luminal patients from an independent medical center (Harbin Medical University Cancer Hospital) thousands of kilometers away from Fujian Medical University Union Hospital (Fig. 3a). Thus, plausible differences in CT and ET from different medical centers did not affect the general trend of the common drug-nonresponsive phenotype. We further tested WTP in 69 TT-treated HER2+ patients from both hospitals (Fig. 1a). In this case, we identified 14 of 22 patients with <5-year DFS against 4 false positives (Fig. 3a), i.e. attained 64% sensitivity and 91% specificity (*p* < 0.001). As a summary, the CT-treated TNBC patients, ET-treated HER2-&luminal patients, and TT-treated HER2+ patients (2) shared a convergent phenotype nonresponsive to anthracycline-taxane, tamoxifen, and trastuzumab, respectively.

### Nonresponse to small molecule drugs among more subtle patient subgroups

The 69 TT-treated HER2+ patients might not represent HER2+ breast cancer well because most HER2+ patients forwent this relatively expensive therapy based on the large molecule drug of trastuzumab antibody, which contrasts sharply with the small molecule drugs of anthracycline-taxane (CT) and tamoxifen (ET). A total of 184 (or 56) HER2+ patients at Fujian Medical University Union Hospital (or Harbin Medical University Cancer Hospital) were offered CT or ET (rather than TT) as their first-line therapy (Fig. 1a), and should be more representative of HER2+ breast cancer. These two patient cohorts exhibited a similar LSHS profile to that of the corresponding CT-treated TNBC patients (Fig. 3a, solid arrows). We will thus limit all discussion below to small molecule drug nonresponse.

Combining the 624 and 213 patients from the two hospitals examined for small molecule drug nonresponse (Fig. 1a), we obtained the corresponding profiles for 5 molecular subtype-classified subgroups by immunohistochemistry (Fig. 3b). Within the luminal patients, there existed a spectrum of profile from luminal A (HSLS) to HER2+&luminal B patients (LSHS), with the profile of HER2-&luminal B patients approximated HSLS more than LSHS. Thus, with common CT treatment, the HER2+&luminal B patients resembled HER2-enriched patients more than other luminal (luminal A and HER2-&luminal B) patients, even though all luminal patients underwent first-line ET in addition to the CT. In this sense, HER2 is a more fundamental molecular marker than luminal (or ER) to stratify patients for personalized therapies (2), rather than the other way around (3). This result strikingly echoed an early report that attributed the poor prognosis of the HER2+&luminal B patients (27) to their strong resistance to tamoxifen (28), and would not be attainable had the HER2-&luminal B patients been treated with TT and trastuzumab (factors that would confound direct comparison of the three luminal subgroups).

Thus, the simplest subgrouping of the 624 patients was to combine luminal A and HER2-&luminal B patients into a HSLS subgroup of HER2-&ER+ patients, and HER2+&luminal B, HER2-enriched, and TNBC patients into a LSHS subgroup of non-HER2-&ER+ patients, except for 2 rare luminal patients that were not ER+ (Fig. 3c). In this way, only 2 immunohistochemistry markers (HER2 and ER) were needed to represent the characteristic dual-profile (HSLS versus LSHS) drug nonresponse.

### Effects of clinical endpoint, cancer progression, and intratumor heterogeneity

To investigate the possible confounding factor of DFS as an imperfect surrogate clinical endpoint for OS (29), we tested WTP on the 624 patients using 5-year OS as the clinical endpoint (Figs. 3a, 3c; broken arrows). In comparison to the prediction on 5-year DFS rate, the prediction on 5-year OS rate reduced false negatives at the cost of increased false positives, due to the imbalanced positive rates of <5-year DFS, <5-year OS, and WTP prediction (Fig. 3c, Inset i). Remarkably, the accuracy to predict 5-year OS rate (75.3%) approximated that on 5-year DFS rate (76.0%), while the characteristic dual-profile drug nonresponse was retained (Fig. 3c). Thus, the prediction of drug nonresponse by this marker was rather independent on the choice of clinical endpoint (DFS versus OS).

Interestingly, the “adaptive” predictive ability of WTP to two clinical endpoints could be extended to two categories of primary tumor size (Figs. 3c, 3d; broken arrows), and to a less degree, to two categories of lymph node status and histological grade (Figs. 3d, right panels). Thus, the prediction of drug nonresponse by this marker was rather independent on tumor progression. We then performed the strictest test on 113 clinicopathologically low-risk patients (among the 624 patients) with small (≤2 cm) tumor size, negative lymph node status, and low histological grade (HG1/HG2) (Fig. 3e; Table S4, early stage). We identified 12 of 18 patients with <5-year DFS against 13 false positives (Fig. 3e), i.e. attained 66% sensitivity and 86% specificity (*p* = 5×10^−5^). This high performance may be beneficial for the risk assessment of screening-detected early stage cancer in a country with breast cancer screening program (16).

It should be noted that our sampling of one FFPE-block per tumor/patient was enough for small (≤2 cm) tumors, but insufficient for large (>2 cm) tumors which might be confounded by block-scale intratumor heterogenicity (Fig. 1b). However, the predictive ability of WTP was rather independent on the two tumor sizes (Fig. 3d, broken arrow), i.e. robust against this intratumor heterogenicity. This feature is highly beneficial for potential clinical translation. Although the block-scale sampling of tumor periphery was robust against the intratumor heterogenicity, the niche-scale imaging revealed both “wound-like” and “non-wound” morphologies (Fig. 2b) in 41% of the patients, in contrast to 52% patients with exclusive “wound-like” or “non-wound” morphology (Fig. 2c) and 7% patients with neither morphology. This 41% portion of patients would be prone to niche-scale intratumor heterogenicity if WTP assessment was not averaged over 5-15 representative tumor peripheral niches (see Methods).

### Prognostic value in the context of routine tumor markers

The above tests of WTP demonstrates its negative predictive value on a convergent drug nonresponse, in contrast to similar uniparameter marker of ER (or HER2) with established positive predictive value on tamoxifen (or trastuzumab) response. This negative predictive value should be differentiated from that on nonbeneficial anthracycline-taxane response, which is often predicted by a multiparameter marker of gene expression assay (18). Because ER and HER2 also have prognostic value (30) just like some routine uniparameter clinicopathological markers (tumor size, histological grade, and lymph node status) (31), it is important to put WTP in the context of all these uniparameter markers and compare their performance to predict prognosis.

The prediction of the 624 patients by 6 uniparameter markers revealed the diverged Kaplan-Meier DFS curves of patients stratified into low- and high-risk groups (Figs. 4a, 4b). The histological grade, ER/HER2, and WTP represented respectively the standard histology, immunohistochemistry, and multiphoton histology, allowing direct comparison of the three methods based on one common FFPE sectioning platform (Fig. 1c). The lymph node status (surgery) and tumor size (gross examination) were from non-microscopy methods (surgery). The WTP marker outperformed all other markers due to a more diverged stratification of patients (higher hazard ratio HR) in the Kaplan-Meier curve with a lower *p* value (higher statistical significance), and a higher accuracy to predict 5-year DFS/OS rate (Figs. 4a, 4b; Table S5). For binary patient subgrouping by each of these markers, WTP further stratified the low- and high-risk subgroups with comparable values of HR and other performance metrics, highlighting its general applicability across different patient subgroups (Fig. 4c, Table S4). The little dependence of the performance on tumor size confirmed robust prognosis against intratumor heterogeneity (Fig. 4c).

**Figure 4.**
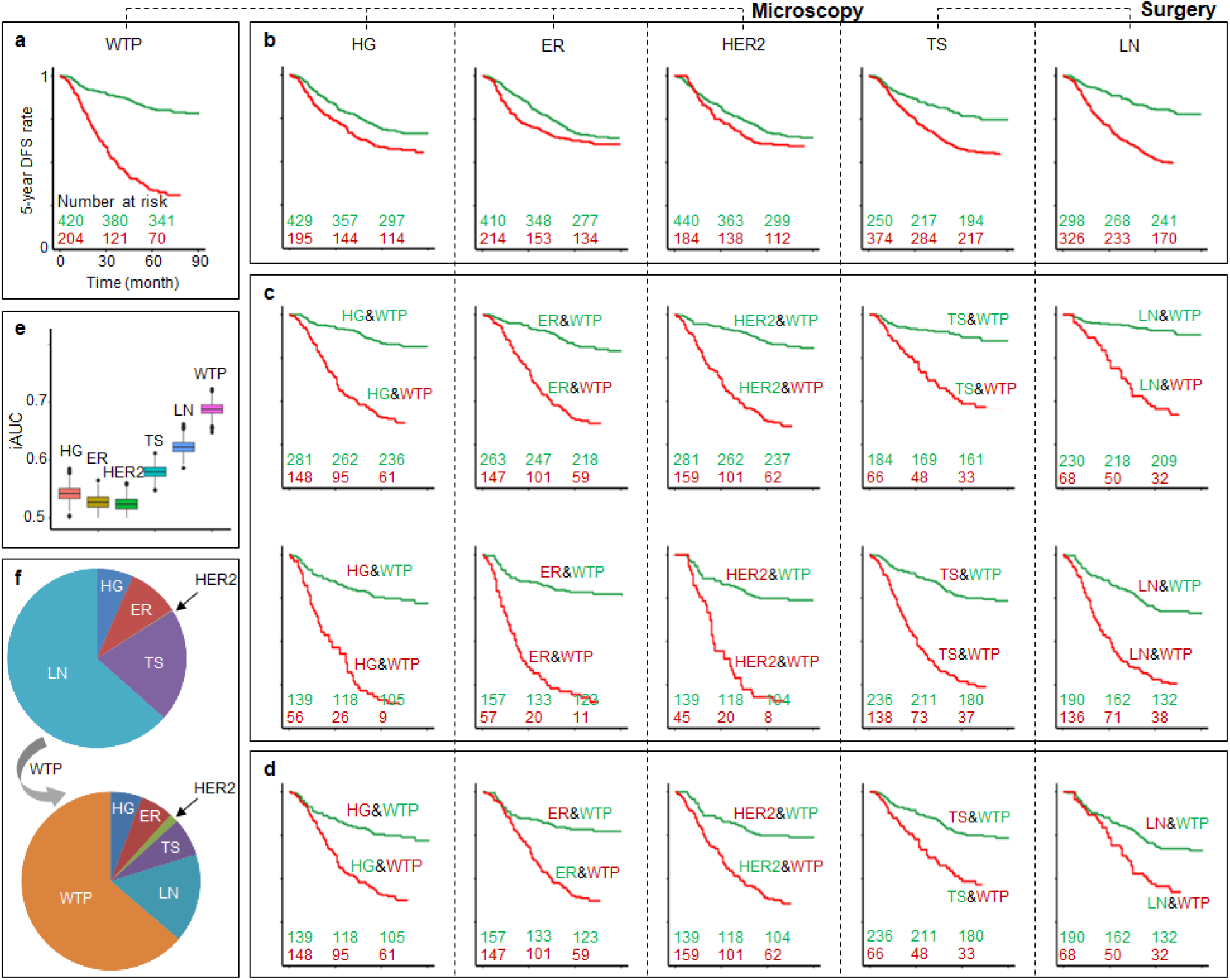
Disease-free survival (DFS) prognosis of 624 patients by WTP. (a) Kaplan-Meier curves according to WTP with numbers of patients at high- and low-risk (red- or green-highlighted numbers, respectively). (b) Kaplan-Meier curves from 5 existing prognostic markers including histological grade (HG), estrogen receptor (ER), human epidermal growth factor receptor 2 (HER2), tumor size (TS), and lymph node status (LN). (c) Kaplan-Meier curves according to WTP for either low-(top) or high-risk patients (bottom) stratified by the 5 markers. (d) Kaplan-Meier curves according to WTP to reclassify significant numbers of low- and high-risk patients classified by the 5 markers. (e) Comparison of prediction accuracy (iAUC) for 6 uniparameter prognostic markers. (f) Relative importance of the 6 markers using multivariate Cox regression analysis (*χ*2 proportion test) for two multiparameter models without/with WTP.

For a significant portion of patients, WTP reclassified the otherwise low-risk (small tumor size, negative lymph node status, low histological grade, ER+, or HER2-) patients to yield a poorer prognosis than their “high-risk” counterparts (large tumor size, positive lymph node status, high histological grade, ER-, or HER2+) (Fig. 4d). Also, WTP achieved the highest incremental area under curve (iAUC), i.e. the most accurate DFS prediction, among the 6 markers (Fig. 4e). Moreover, for risk stratification capability (high HR and low *p* value), the WTP-only prognosis outperformed or approximated various multiparameter regression models that combined all routine prognostic markers of breast cancer (Table S5). Finally, the relative importance of prognostic markers using multivariate Cox regression analysis (Table S6) indicated the dominant contribution of WTP to DFS (Fig. 4f). Similar results were obtained from the OS prognosis of these patients (Fig. S4, Tables S4-S6). Since all 5 contextual prognostic markers have a long history of development and optimization, it is surprising to witness the emergence of a dominant prognostic marker like WTP (the underlying mechanism of which may have long been missed in routine breast cancer prognosis).

## Discussion

The observed dual-profile (HSLS versus LSHS) drug nonresponse consists of four patient subgroups along two dimensions (Fig. 3c, Inset ii), who account for 76% of the patient population at Fujian Medical University Union Hospital and 80% of the patient population at Harbin Medical University Cancer Hospital (Fig. 3a). Along the molecular dimension, HER2-&ER+ patients predictive of tamoxifen response can be separated from non-HER2-&ER+ patients predictive of anthracycline-taxane response. Along the morphological dimension, WTP+ patients predictive of the convergent drug nonresponse can be separated from WTP-patients predictive of good prognosis (Fig. 3c, Inset ii; Fig. 4a). This pattern of drug nonresponse is highly profound and not obscured by covariate noises such as hosting hospital, clinical endpoint, tumor size, lymph node status, histological grade, and patient age. Thus, often-overlooked morphological markers may be as important as the best molecular markers, and microscopy should remain a cornerstone of surgical pathology (32). Our molecular subgrouping of HER2-&ER+ versus non-HER2-&ER+ patients reproduces the profound luminal versus basal (myoepithelial) subgrouping by intrinsic gene subset cluster analysis (1). The large number of false positives for HER2-&ER+ patients (Fig. 3c) may be due to the late recurring dynamics (or cancer dormancy) unique to the ER+ patients (17). Extended follow-up from 5-year DFS/OS rate in this study to 10-year DFS/OS rate in the future will test this hypothesis and may expand the applicability of the characteristic dual-profile pattern. The remaining critical deficiency of WTP, i.e. the large extent of false negatives for non-HER2-&ER+ patients (Fig. 3c), requires a molecular interpretation just like how the morphological marker of histological grade is interpreted based on the signaling pathways of cell cycle regulation and proliferation (33). Assuming independent patient subgrouping along the two dimensions (Fig. 3c, Inset ii), the underlying molecular mechanism of WTP should be independent of breast cancer molecular subtyping, i.e. be a pan-cancer molecular mechanism.

The apparent link of WTP with wound-like tumor peripheral niches (34) supporting migratory (mesenchymal) tumor cells and multidrug resistance (4-6) suggests an underlying mechanism of EMT, which is a well-known pan-cancer molecular mechanism (26). More lines of indirect evidence support this interpretation. First, circulating tumor cells exhibit (partial) EMT and drug resistance in luminal, HER2+, and TNBC patients (35) just like the convergent drug nonresponse exists across these patients (Fig. 3a), while the increased extent of false negatives from HER2-&ER+ to non-HER2-&ER+ patients (Fig. 3c) parallels the increased (EMT-dispensable) collective cell invasion from luminal to TNBC patients (36). Thus, WTP may be a surrogate EMT marker dispensable for metastasis but indispensable for drug resistance (8). Second, the observed WTP+ at the earliest clinicopathological stage (Fig. 3e) parallels the temporal EMT activation early in the multistep carcinoma progression of mice (37) and humans (38). Third, the spatially heterogenous distribution of WTP+ and WTP-niches at primary tumor periphery parallels the enrichment of post-EMT cells at tumor invasion fronts (39) and heterogeneous EMT transition states localized in different niches (40). Fourth, human residual breast cancers resistant to neoadjuvant ET (letrozole) or CT (docetaxel) display mesenchymal and tumor-initiating features correlative with poor prognosis (41), even though the corresponding gene expression marker was linked to a specific type of molecular subtype (“claudin-low”) rather than a pan-cancer marker independent of breast cancer molecular subtyping (42). Despite these lines of indirect evidence, the ultimate confirmation of the EMT nature of WTP will require changes in a set of molecular markers and cellular properties (43) to synchronize with the WTP marker of 2.8-mm niche-scale extracellular matrix remodeling (44).

The role of EMT in metastasis has been gradually clarified in cultured cells and animal models but remained controversial in humans. Pathologists would embrace this role if a prototypical EMT marker could produce large independent prognostic values after adjustment for routine prognostic markers (45). However, EMT-inspired histopathological markers such as tumor budding have not demonstrated a large prognostic value for routine clinical application (46). The uniqueness of WTP lies in a shifted emphasis from tumor cells (standard histology) to extracellular matrix (multiphoton histology), so that overall EMT state may be indirectly assessed from its synchronized process of extracellular matrix remodeling (43). In this way, the challenge to differentiate rare mesenchymal tumor cells among abundant fibroblasts (46) can be avoided. Label-free multiphoton microscopy developed in early 2000s (i.e. multiphoton histology) has excelled in imaging both the extracellular matrix and (tumor) cells in live tissue (47), but its differential benefit over standard histology remains ambiguous due to typical incompatibility of the two approaches. By “avoiding” the often cherished three-dimensional and live-tissue imaging afforded by multiphoton microscopy, our co-registered standard-multiphoton histology synergistically utilizes standard histology to recognize primary tumor periphery and multiphoton histology to detect WTP (Fig. 1d). Surprisingly, this technology demonstrates a large differential value of WTP over histological grade, ER, and HER2 in microscopy-based cancer prognosis. By comparing the co-reregistered images of standard and multiphoton histology (Figs. 2b, 2c), we conclude that the WTP marker can be reliably assessed by the multiphoton histology but not the standard histology. Thus, the methodology established in this study may motivate the clinical translation of multiphoton histology and other novel optical imaging technologies (48) via the blueprint of immunohistochemistry in the 1980s and whole slide imaging more recently (Fig. 1c).

Regardless of the underlying molecular mechanism, WTP from co-registered standard-multiphoton histology achieves both high performance and low complexity for general cancer prognosis, in comparison to closely related alternatives such as uniparameter (11) or multiparameter markers from gene-expression assay (18), tumor infiltrating lymphocytes (49, 50) or lymphovascular invasion (51) from standard histology or immunohistochemistry, and tumor-associated collagen signatures (TACS) from label-free multiphoton microscopy (52) recently integrated with standard histology (53). These alternatives are limited by at least one of the following factors. First, the markers are often correlated with certain endpoints by a pair of “training” and “validation” patient cohorts using adjustable (fitting) variables, which may be susceptible to various covariate noises. In contrast, our uniparameter-WTP prediction employs no fitting variable for both DFS and OS endpoints, and our patients from the two hospitals are simply two independent cohorts to test WTP (because no fitting variables are needed to train and validate a model). It would be challenging to predict both DFS and OS equally well (76% vs. 75% accuracy, see Fig. 3c) without fitting variable(s). Second, incompatibility with standard histology and/or instability against intratumor heterogeneity hinder clinical translation. This limiting factor is prominent for the gene-expression assay and other non-imaging molecular analyses (25), including that developed a uniparameter marker (11), but is absent in our multiphoton imaging assay due to the representative sampling of tumor peripherical niches. Third, high complexity (or cost) prohibits user-friendly application. The (gene-expression) molecular assay is often limited by the high complexity of multiregional sampling to mitigate the intratumor heterogeneity, the standard histology or immunohistochemistry by well-trained pathologists to examine the tumor infiltrating lymphocytes and lymphovascular invasion, and the multiphoton microscopy by imaging experts to identify 8 distinct morphologies of TACS (53), most of which may present as confounding factors to obscure a causal understanding of the underlying mechanism. In contrast, we find that WTP assessment by naive human eyes is fast and consistent (Figs. S1-S3), not only among the three inspectors (see Methods) but also other laypersons after a brief (10-hr) training.

Finally, we comment on possible limitations of this study arising from its “idealized” design. First, our technology is not applicable to nonmetastatic breast cancer patients undergoing neoadjuvant therapy due to the corresponding disruption to their tumor microenvironment. On the other hand, the dominant role of WTP over routine prognostic markers may neutralize one prominent advantage of the neoadjuvant therapy over its adjuvant counterpart, i.e. pCR as short-term endpoint to predict drug response. Second, our patients have been restricted to symptom-detected patients with a typically poorer prognosis than that of screening-detected patients. The applicability of WTP to screening-detected patients is nevertheless promising due to the validity of WTP at the earliest clinicopathological stage. Third, our patients have been restricted to those undergoing CT to detect the convergent drug nonresponse. This is useful for identifying high-risk patients who would not benefit from conventional therapies (CT, ET, and TT) due to their intrinsically poor prognosis, but not necessarily useful for identifying low-risk patients who would not benefit from the CT due to their intrinsically good prognosis. The gene expression assay and computational risk assessment suit these low-risk patients well. Fourth, we note that WTP relies heavily on collagen-rich tumor microenvironment afforded by breast cancer and may not be applicable to collagen-poor cancer types such as liver cancer. However, we believe similar tumor microenvironment-based morphological markers may be developed (as a surrogate for EMT) for these cancer types to innovate cancer histopathology in general.

## Data Availability

All data needed to evaluate the conclusions in this paper are present in the manuscript or the supplementary materials and all collected images are available upon request.

## Acknowledgements

This work was partially supported by grants from the National Institutes of Health, U.S. Department of Health and Human Services (R01 CA241618, R43 MH119979, and R41 GM139528). Also, this work was supported by the Natural Science Foundation of China (Grant No. 82171991, 82172800), Special Funds of the Central Government Guiding Local Science and Technology Development (No. 2020L3008), Fujian Major Scientific and Technological Special Project for “Social Development” (No. 2020YZ016002), Natural Science Foundation of Fujian Province (No. 2019J01269, No. 2020J011008, and No. 2020J01839), and Joint Funds for the Innovation of Science and Technology of Fujian Province (2019Y9101).

