## Supplementary material for "Wound-like tumor periphery in human breast cancer predicts a convergent drug nonresponse": Figures S1-S4 and Tables S1-S6

### Supplementary Materials:

#### Figures S1-S4

#### Tables S1-S6

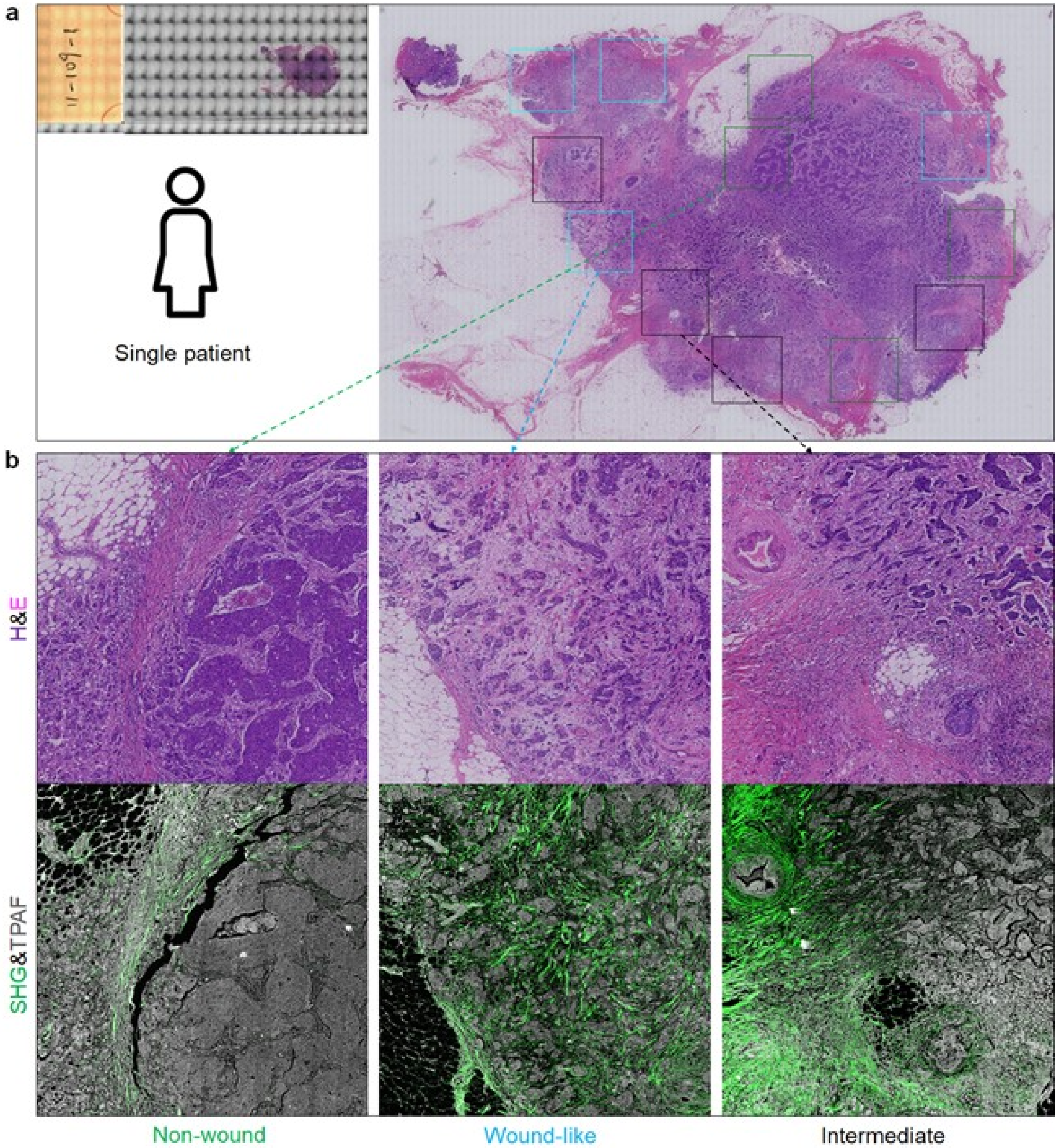


**Fig. S1.** Visualization of two competing tumor peripheries in one patient. Scale of magnified square images: 2.8 mm. (a) H&E-stained microscope slide of the patient and H&E image with multiple niches (marked squares) at the invasive fronts. (b) Observed “non-wound” (green square), “wound-like” (blue square), and “intermediate” (black square) tumor peripheries with co-registered SHG-TPAF (green-gray) and H&E images, in which the “intermediate” state is largely a spatial combination of the “non-wound” and “wound-like” tumor peripheries.

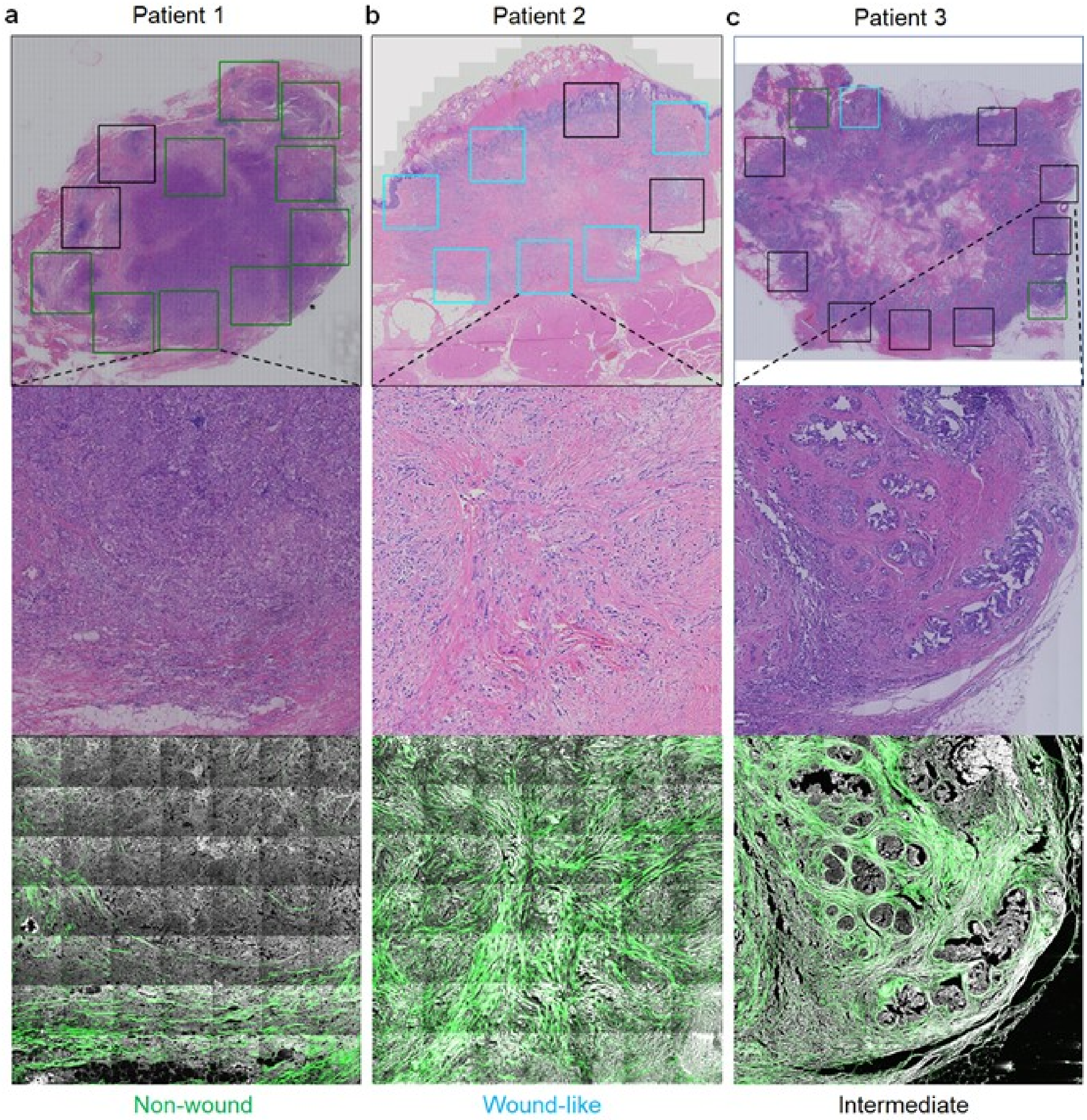


**Fig. S2.** Comparison of two competing tumor peripheries in three patients. Scale of magnified square images: 2.8 mm. (a) H&E and co-registered SHG-TPAF images obtained from one patient dominated by the "non-wound" tumor periphery (green squares). (b) Similar images from another patient dominated by the "wound-like" tumor periphery (blue squares). (c) Similar images from a third patient dominated by the "intermediate" tumor periphery (black squares), in which the "intermediate" state contains some mammary ducts and is largely a functional combination of the "non-wound" and "wound-like" tumor peripheries.

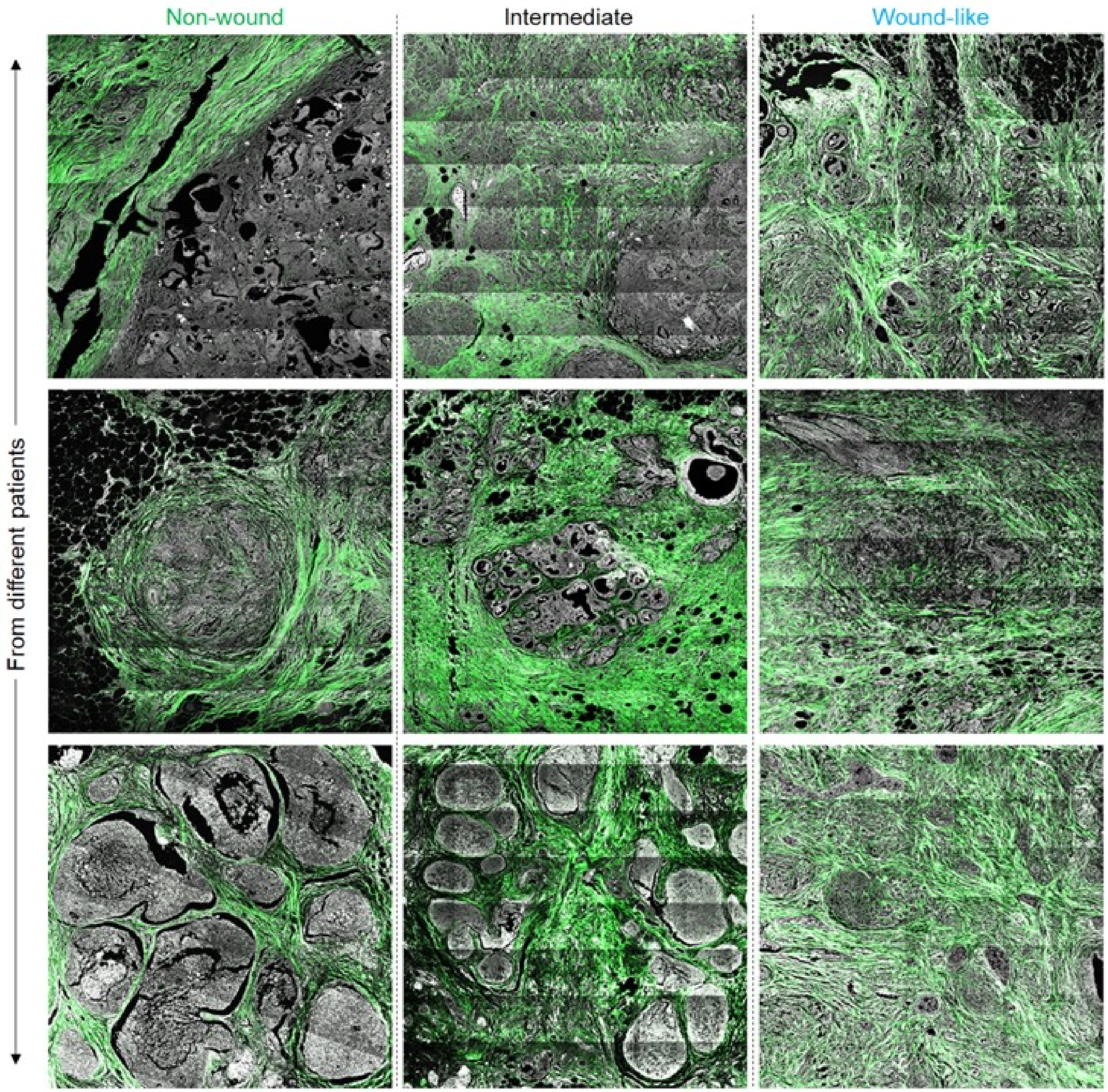


**Fig. S3.** Representative examples of “non-wound”, “wound-like”, and “intermediate” tumor peripheral niches observed from different patients.

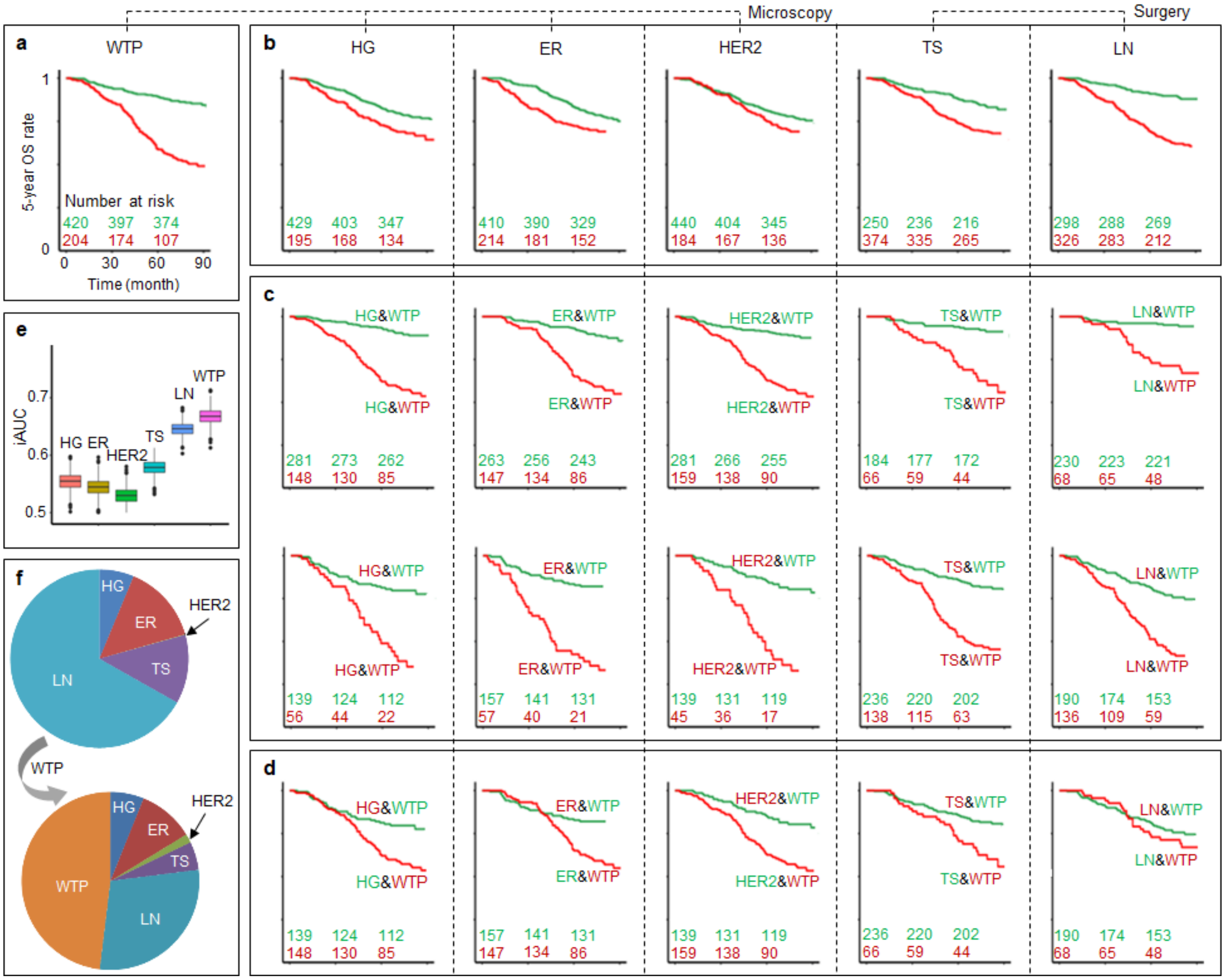


**Fig. S4.** Overall survival (OS) prognosis of 624 patients by WTP. (a) Kaplan-Meier curves according to WTP with numbers of patients at high- and low-risk (red- or green-highlighted numbers, respectively). (b) Kaplan-Meier curves from 5 existing prognostic markers including histological grade (HG), estrogen receptor (ER), human epidermal growth factor receptor 2 (HER2), tumor size (TS), and lymph node status (LN). (c) Kaplan-Meier curves according to WTP for either low- (top) or high-risk patients (bottom) stratified by the 5 markers. (d) Kaplan-Meier curves according to WTP to reclassify significant numbers of low- and high-risk patients classified by the 5 markers. (e) Comparison of prediction accuracy (iAUC) for 6 uniparameter prognostic markers. (f) Relative importance of the 6 markers using the Cox regression analysis ($\chi^2$ proportion test) for two multiparameter models without/with WTP.

**Table S1.** Symptom-detected breast cancer patients from two medical centers.

| Characteristics | Fuzhou patients undergoing CT (n = 624) | Harbin patients undergoing CT (n = 213) |
|---|---|---|
| 5-yr DFS rate | 408 (65.4%) | 134 (62.9%) |
| 5-yr OS rate | 478 (76.6%) | |
| **Age** | | |
| ≤50 | 365 (58.5%) | 115 (54.0%) |
| >50 | 259 (41.5%) | 98 (46.0%) |
| **Molecular subtype** | | |
| Luminal A | 133 (21.3%) | 55 (25.8%) |
| HER2-&luminal B | 202 (32.4%) | 64 (30.0%) |
| HER2+&luminal B | 78 (12.5%) | 26 (12.2%) |
| HER2 enriched | 106 (17.0%) | 30 (14.1%) |
| TNBC | 105 (16.8%) | 38 (17.8%) |
| **Tumor size** | | |
| ≤2cm | 250 (40.0%) | 112 (52.6%) |
| 2-5cm | 332 (53.2%) | 95 (44.6%) |
| >5cm | 42 (6.7%) | 6 (2.8%) |
| **Lymph node status** | | |
| 0 | 298 (47.8%) | 89 (41.8%) |
| 1-3 | 154 (24.7%) | 69 (32.4%) |
| ≥4 | 172 (27.6%) | 55 (25.8%) |
| **Histological grade** | | |
| HG1 | 99 (15.9%) | 6 (2.8%) |
| HG2 | 330 (52.9%) | 172 (80.8%) |
| HG3 | 195 (31.3%) | 35 (16.4%) |

**Table S2.** Age-related characteristics of 624 patients from main medical center.

| Characteristics | Fuzhou luminal patients (n = 413) | Fuzhou non-luminal patients (n = 211) |
|---|---|---|
| 5-yr DFS rate | 274 (66.3%) | 134 (63.5%) |
| 5-yr OS rate | 327 (79.2%) | 151 (71.6%) |
| Age ≤50 | 252 (61.0%) | 113 (53.6%) |
| Age >50 | 161 (39.0%) | 98 (46.4%) |
| Average age | 48.0 | 49.5 |

**Table S3.** Distributions of WTP1-3 in the 624 patients subdivided binarily by 5 routine prognostic markers.

| Characteristics | Total (624) | WTP1 (348) | WTP2 (72) | WTP3 (204) |
|---|---|---|---|---|
| 5-yr DFS rate | 408 (65.4%) | 289 (83.0%) | 50 (69.4%) | 69 (33.8%) |
| 5-yr OS rate | 478 (76.6%) | 315 (90.5%) | 57 (79.2%) | 106 (52.0%) |
| **ER** | | | | |
| positive | 410 (65.7%) | 215 (61.8%) | 48 (66.7%) | 147 (72.1%) |
| negative | 214 (34.3%) | 133 (38.2%) | 24 (33.3%) | 57 (27.9%) |
| **HER2** | | | | |
| positive | 184 (29.5%) | 113 (32.5%) | 26 (36.1%) | 45 (22.1%) |
| negative | 440 (70.5%) | 235 (67.5%) | 46 (63.9%) | 159 (77.9%) |
| **Histological grade** | | | | |
| HG1/HG2 | 429 (68.8%) | 227 (65.2%) | 54 (75.0%) | 148 (72.5%) |
| HG3 | 195 (31.2%) | 121 (34.8%) | 18 (25.0%) | 56 (27.5%) |
| **Tumor size** | | | | |
| ≤2cm | 250 (40.1%) | 146 (42.0%) | 38 (52.8%) | 66 (32.4%) |
| >2cm | 374 (59.9%) | 202 (58.0%) | 34 (47.2%) | 138 (67.6%) |
| **Lymph node status** | | | | |
| positive | 326 (52.2%) | 153 (44.0%) | 37 (51.4%) | 136 (66.7%) |
| negative | 298 (47.8%) | 195 (56.0%) | 35 (48.6%) | 68 (33.3%) |

**Table S4.** Prediction of DFS/OS prognosis of the 624 patients by WTP in different subgroups.

| | 5-yr **DFS** | WTP- | HR | 95% CI | *p* | Sensitivity | Specificity | Accuracy |
|---|---|---|---|---|---|---|---|---|
| *Treatment group* | | | | | | | | |
| TNBC | 62 (59.0%) | 74 (70.5%) | 7.84 | 4.20-14.64 | 1E-10 | 0.628 | 0.935 | 0.810 |
| HER2-&luminal | 234 (69.9%) | 207 (61.8%) | 4.84 | 3.23-7.25 | 2E-14 | 0.703 | 0.756 | 0.740 |
| HER2+ | 112 (60.9%) | 139 (75.5%) | 5.28 | 3.32-8.41 | 2E-12 | 0.514 | 0.929 | 0.766 |
| *Early stage* | 95 (84.1%) | 88 (77.9%) | 6.40 | 2.61-15.70 | 5E-05 | 0.667 | 0.863 | 0.832 |
| *Tumor size* | | | | | | | | |
| ≤2cm | 192 (76.8%) | 184 (73.6%) | 4.66 | 2.81-7.72 | 2E-09 | 0.586 | 0.833 | 0.776 |
| >2cm | 216 (57.8%) | 236 (63.1%) | 4.70 | 3.42-6.46 | 2E-21 | 0.639 | 0.829 | 0.749 |
| *Lymph node status* | | | | | | | | |
| negative | 240 (80.5%) | 230 (77.2%) | 6.76 | 4.10-11.14 | 6E-14 | 0.638 | 0.871 | 0.826 |
| positive | 168 (51.5%) | 190 (58.3%) | 3.43 | 2.49-4.71 | 3E-14 | 0.620 | 0.774 | 0.699 |
| *Histological grade* | | | | | | | | |
| HG1/HG2 | 294 (68.5%) | 281 (65.5%) | 4.83 | 3.43-6.80 | 1E-19 | 0.652 | 0.796 | 0.751 |
| HG3 | 114 (58.5%) | 139 (71.3%) | 5.51 | 3.55-8.55 | 3E-14 | 0.580 | 0.921 | 0.779 |
| *ER* | | | | | | | | |
| positive | 274 (66.8%) | 263 (64.1%) | 4.50 | 3.20-6.32 | 5E-18 | 0.654 | 0.788 | 0.744 |
| negative | 134 (62.6%) | 157 (73.4%) | 6.32 | 4.06-9.82 | 3E-16 | 0.575 | 0.918 | 0.790 |
| *HER2* | | | | | | | | |
| positive | 112 (60.9%) | 139 (75.5%) | 5.28 | 3.32-8.41 | 2E-12 | 0.514 | 0.929 | 0.766 |
| negative | 296 (67.3%) | 281 (63.9%) | 5.11 | 3.65-7.15 | 2E-21 | 0.681 | 0.794 | 0.757 |
| | 5-yr **OS** | WTP- | HR | 95% CI | *p* | Sensitivity | Specificity | Accuracy |
| *Treatment group* | | | | | | | | |
| TNBC | 70 (66.7%) | 74 (70.5%) | 5.47 | 2.72-11.02 | 2E-06 | 0.629 | 0.871 | 0.790 |
| HER2-&luminal | 273 (81.5%) | 207 (61.8%) | 4.20 | 2.59-6.97 | 3E-08 | 0.774 | 0.707 | 0.719 |
| HER2+ | 135 (73.4%) | 139 (75.5%) | 4.41 | 2.60-7.49 | 4E-08 | 0.571 | 0.874 | 0.793 |
| *Early stage* | 102 (90.3%) | 88 (77.9%) | 6.31 | 2.06-19.30 | 1E-03 | 0.727 | 0.833 | 0.823 |
| *Tumor size* | | | | | | | | |
| ≤2cm | 214 (85.6%) | 184 (73.6%) | 4.91 | 2.65-9.11 | 4E-07 | 0.639 | 0.799 | 0.776 |
| >2cm | 264 (70.6%) | 236 (63.1%) | 3.46 | 2.38-5.02 | 7E-11 | 0.682 | 0.761 | 0.738 |
| *Lymph node status* | | | | | | | | |
| negative | 267 (89.6%) | 230 (77.2%) | 5.97 | 2.98-11.92 | 4E-07 | 0.677 | 0.824 | 0.809 |
| positive | 211 (64.7%) | 190 (58.3%) | 2.85 | 1.99-4.09 | 1E-08 | 0.670 | 0.720 | 0.702 |
| *Histological grade* | | | | | | | | |
| HG1/HG2 | 345 (80.4%) | 281 (65.5%) | 4.68 | 3.07-7.12 | 7E-13 | 0.762 | 0.757 | 0.758 |
| HG3 | 133 (68.2%) | 139 (71.3%) | 3.53 | 2.14-5.83 | 8E-07 | 0.548 | 0.835 | 0.744 |
| *ER* | | | | | | | | |
| positive | 327 (79.8%) | 263 (64.1%) | 3.91 | 2.57-5.93 | 2E-10 | 0.747 | 0.740 | 0.741 |
| negative | 151 (70.6%) | 157 (73.4%) | 4.85 | 2.95-7.98 | 5E-10 | 0.571 | 0.861 | 0.776 |
| *HER2* | | | | | | | | |
| positive | 135 (73.4%) | 139 (75.5%) | 4.41 | 2.60-7.49 | 4E-08 | 0.571 | 0.874 | 0.793 |
| negative | 343 (78.0%) | 281 (63.9%) | 4.23 | 2.81-6.36 | 5E-12 | 0.722 | 0.741 | 0.736 |

**Table S5.** Prediction of 5-year DFS/OS prognosis of the 624 patients by different uniparameter markers and multiparameter regression models.

| Biomarkers for observed <5-year DFS rate (34.6%) | Predicted high risk | HR | *p* | Sensitivity | Specificity | Accuracy |
|---|---|---|---|---|---|---|
| TS (>2 cm vs ≤2 cm) | 374 (59.9%) | 2.060 | 1E-06 | 0.731 | 0.471 | 0.561 |
| LN (+ vs. -) | 326 (52.2%) | 2.861 | 8E-13 | 0.731 | 0.588 | 0.638 |
| HG (HG3 vs. HG1/HG2) | 195 (31.2%) | 1.450 | 0.007 | 0.375 | 0.721 | 0.601 |
| ER (- vs. +) | 214 (34.3%) | 1.243 | 0.113 | 0.370 | 0.672 | 0.567 |
| HER2 (+ vs. -) | 184 (29.5%) | 1.240 | 0.127 | 0.333 | 0.725 | 0.590 |
| WTP (+ vs. -) | 204 (32.7%) | 4.831 | 1E-30 | 0.625 | 0.831 | 0.760 |
| TS, LN, HG, ER, HER2 (2-category, see above) | 258 (41.3%) | 3.097 | 1E-16 | 0.639 | 0.706 | 0.683 |
| TS, LN, HG, MS (>2-category, see Note)* | 145 (23.2%) | 4.710 | 7E-31 | 0.481 | 0.900 | 0.755 |
| Biomarkers for observed 5-year OS rate (76.6%) | | | | | | |
| TS (>2 cm vs ≤2 cm) | 374 (59.9%) | 2.054 | 6E-05 | 0.753 | 0.448 | 0.519 |
| LN (+ vs. -) | 326 (52.2%) | 3.975 | 1E-12 | 0.788 | 0.559 | 0.612 |
| HG (HG3 vs. HG1/HG2) | 195 (31.2%) | 1.594 | 0.004 | 0.425 | 0.722 | 0.652 |
| ER (- vs. +) | 214 (34.3%) | 1.445 | 0.023 | 0.432 | 0.684 | 0.625 |
| HER2 (+ vs. -) | 184 (29.5%) | 1.31 | 0.105 | 0.336 | 0.718 | 0.628 |
| WTP (+ vs. -) | 204 (32.7%) | 3.990 | 2E-17 | 0.671 | 0.778 | 0.780 |
| TS, LN, HG, ER, HER2 (2-category, see above) | 258 (41.3%) | 3.918 | 2E-15 | 0.712 | 0.678 | 0.686 |
| TS, LN, HG, MS (>2-category, see Note)* | 146 (23.4%) | 5.452 | 3E-26 | 0.575 | 0.870 | 0.770 |

*Note: TS - tumor size (≤2 cm, 2-5 cm, and >5 cm); LN - lymph node status (0, 1-3, and ≥4); HG - histological grade (HG1, HG2, and HG3); and MS - Molecular subtype (Luminal A, HER2-&luminal B, HER2+&luminal B, HER2 enriched, TNBC)

**Table S6.** Multivariate Cox proportional hazards regression analysis of the association of 6 markers with 5-year DFS/OS rate of the 624 patients.

| Low-risk | Patients | 5-year **DFS** | Multivariate analysis | | | |
|---|---|---|---|---|---|---|
| High-risk | 624 | 408 (65.4%) | HR | (95%CI) | | *p* |
| TS: ≤2 cm | 250 | 192 (76.8%) | Reference | | | |
| TS: >2 cm | 374 | 216 (57.8%) | 1.643 | 1.224 | 2.206 | 0.001 |
| LN: - | 298 | 240 (80.5%) | Reference | | | |
| LN: + | 326 | 168 (51.5%) | 2.273 | 1.683 | 3.069 | 8E-08 |
| HG1/HG2 | 429 | 294 (68.5%) | Reference | | | |
| HG3 | 195 | 114 (585%) | 1.512 | 1.148 | 1.990 | 0.003 |
| ER+ | 410 | 274 (66.8%) | Reference | | | |
| ER- | 214 | 134 (62.6%) | 1.619 | 1.188 | 2.205 | 0.002 |
| HER2- | 440 | 296 (67.3%) | Reference | | | |
| HER2+ | 184 | 112 (60.9%) | 1.227 | 0.900 | 1.673 | 0.195 |
| WTP- | 420 | 339 (80.7%) | Reference | | | |
| WTP+ | 204 | 69 (33.8%) | 4.908 | 3.708 | 6.497 | 9E-29 |
| Low-risk | Patients | 5-year **OS** | | | | |
| High-risk | 624 | 478 (76.6%) | HR | (95%CI) | | *p* |
| TS: ≤2 cm | 250 | 214 (85.6%) | Reference | | | |
| TS: >2 cm | 374 | 264 (70.6%) | 1.489 | 1.041 | 2.130 | 0.029 |
| LN: - | 298 | 267 (89.6%) | Reference | | | |
| LN: + | 326 | 211 (64.7%) | 3.300 | 2.232 | 4.878 | 2E-09 |
| HG1/HG2 | 429 | 345 (80.4%) | Reference | | | |
| HG3 | 195 | 133 (68.2%) | 1.497 | 1.081 | 2.072 | 0.015 |
| ER+ | 410 | 327 (79.8%) | Reference | | | |
| ER- | 214 | 151 (70.6%) | 1.827 | 1.280 | 2.606 | 0.001 |
| HER2- | 440 | 343 (78.0%) | Reference | | | |
| HER2+ | 184 | 135 (73.4%) | 1.194 | 0.833 | 1.711 | 0.335 |
| WTP- | 420 | 372 (88.6%) | Reference | | | |
| WTP+ | 204 | 106 (52.0%) | 3.749 | 2.696 | 5.214 | 4E-15 |

Note: TS - tumor size; LN - lymph node status; HG - histological grade.
